# Bacteriophages can be used as infection control agents: A proof-of concept study involving anti-*Acinetobacter baumannii* phage

**DOI:** 10.1101/852582

**Authors:** Aamir Hussain, Amna Manzoor, Ihsan Ullah, Atif Aziz, Mubashar Aziz, Muhammad Qamar Saeed

## Abstract

Hospital acquired infections are responsible for morbidity and mortality worldwide. *Acinetobacter spp*. infections are particularly notorious for complicating patient management in ICU settings. Extremely high mortality rates are associated with *Acinetobacter* infections because of their resistance to first- and second-line drugs. There is imminent need to develop infections control systems that are specific and environment friendly. Here, we report a proof-of-concept anti-*Acinetobacter spp*. bacteriophage-based infection control assay that is very target specific as well as innocuous to environment. extensively drug resistant (XDR) *Acinetobacter baumannii* strain was inoculated at various solid surfaces. A bacteriophage, enriched in the same strain, was applied on the inoculated surfaces. Phenol (carbolic acid) was used as a positive control. We show that bacteriophages can be used as infection control agents. In our assay, they killed XDR *Acinetobacter baumannii* present on solid surfaces. Our bacteriophage was extremely effective at reducing the CFU of inoculated strain to almost undetectable levels.

## Introduction

Different surfaces in healthcare facilities may serve as reserves of pathogens. These pathogens are originally contributed by infected hospitalized patients [1]. Bacteria are well known to be promiscuous in terms of exchange of genetic material. They can donate or accept genes even from distantly related species and strains directly or indirectly [2,3]. Under selective evolutionary pressures in hospitals, these pathogens, having genetic capability of horizontal gene transfer, gradually evolve increased virulence and antibiotic resistance— the characteristic of hospital acquired infections [2]. Critical first step of infection control, therefore, is providing aseptic environment to patients so that cost of treatment as well as transmission of multidrug resistant (MDR) infections can be avoided.

There are many routes of transmission of infection in hospital settings. Direct contact between patients is a rare occurrence. A health care worker who may carry some pathogen on his/her skin or clothing may be the source of infection [1,4]. Similarly, patients are in direct contact with ward and ICU surfaces. Pathogens can also hide in medical devices, which, when applied to patients, can transmit infections [1]. MDR *Pseudomonas aeruginosa*, *Staphylococcus aureus* and *Acinetobacter baumannii* infections are mostly acquired in hospital settings [5].

Clinically isolated strains of *Acinetobacter spp*. are particular notorious for their extreme drug resistance. This organism has the ability to cause different systemic infections which include bacteremia, pneumonia, urinary tract infections, superficial and deep wound infections, and meningitis etc. [3]. Ability to cause multiple types of infections, coupled with drug resistance greatly limits treatment options. The best options against it is to develop targeted infection control regimes which are effective at clearing *Acinetobacter spp.* from various surfaces acting as their reservoirs.

Many hospitals rely on carbolic acid (phenol) as their method of disinfection. However, phenolics are damaging to both environment and humans [6,7]. These compounds may cause skin corrosion, germ cell mutagenicity and organ toxicity on repeated exposures. Moreover, use of phenolics in nurseries is highly discouraged. In one study, phenolics exposed infants were found to have higher bilirubin levels as compared to non-exposed, even when disinfectant was prepared according to manufacturer’s recommendation [6]. Therefore, such methods must not be used in medical instruments such as ventilators. There is need of bactericidal agents that are effective as well as non-toxic to patients.

Bacteriophages are natural bacterial parasites. They have the benefit of being extremely specific to their host species and are innocuous to environment and humans [8]. Additionally, as they reproduce in their host, they keep on growing on the surfaces where their host is available. Thirdly, they are self-limiting. They survive only as long as their host survives [9]. Use of bacteriophages in critical health care instrument, therefore, can be very useful in controlling infections in health care settings.

We designed a proof-of-concept study to show that bacteriophage raised against *XDR Acinetobacter baumannii* could effectively reduce bacterial load on solid surfaces. Our assay shows that it was almost as effective as phenol in reducing the colony forming units on inoculated surfaces.

## Materials and Methods

### Selection of Acinetobacter spp. for Phage Isolation

A patient isolated strain of *Acinetobacter baumannii* was selected because it was resistant to all the tested drugs. This strain was provided by Armed Forces Institute of Pathology, Rawalpindi. This strain was used to isolate bacteriophage from sewage samples.

*Acinetobacter baumannii* was cultured on Blood and MacConkey agar plates a day before the start of bacteriophage isolation. For the isolation of bacteriophages, sewage water samples were collected from multiple sites in sterile pre-labeled bottles. Sampling sites included effluents from combined military hospital (CMH) Multan, Nishtar Hospital Multan, Children Hospital Multan and some dairy farms effluent from Multan region.

### Phage enrichment

A heavy inoculum of sub cultured *Acinetobacter baumannii* was prepared from a randomly selected colony on MacConkey agar in a 10 ml tube by overnight culture. Pre-labeled glass flasks were poured with 10 ml of 2x Luria-Bertani (LB) broth (Oxoid, Catalog no. 0264) followed by addition of 10 ml sterile sewage samples filtrate and 500 µl of bacterial culture. Sewage samples were sterilized using 0.2 μl syringe filters (Corning, Catalog no. 411224). All these flasks were incubated at 37⁰C for 24 hours.

After 24 hours, 12-13 ml suspension from each flask was transferred into pre labelled 15 ml tubes. After centrifugation of 10 minutes at 4200 RPM, the supernatant from each tube was syringe filtered. This filtrate was used for two more rounds of enrichment to increase titers of possible anti-acinetoviruses.

### Plaque assay

After three enrichments, suspected bacteriophage containing filtrates were taken and serially diluted ten times (10^−1^ through 10^−10^) in sterile saline solution in ten tubes. An early log phase bacterial inoculum was prepared by adding 300 μl of bacterial suspension in 30 ml LB broth in a flask, followed by the incubating it for 2-4 hours at 37°C. After that, a mixture of bacterial isolate, bacteriophage dilution and soft LB agar (normal solid media contain 1.5% agar) was prepared by adding 0.6% agar in LB broth. For this purpose, 100 μl of early log phase inoculum was added into 20 ml glass tube followed by the addition of 10μl of each of the ten phage dilutions. Then 20 ml of soft LB agar was added. All these tubes were incubated at 37⁰C for 20 minutes for adsorption to take place.

After that about 8 ml of the mixture from each tube was poured on pre labeled LB agar plates and incubated at 37⁰C for 24 hours or until appearance of plaques on phage positive plates. Plaques were observed and virus titer was calculated by counting plaques on 10^−7^ plate (Fig 1).

**Fig 1:**
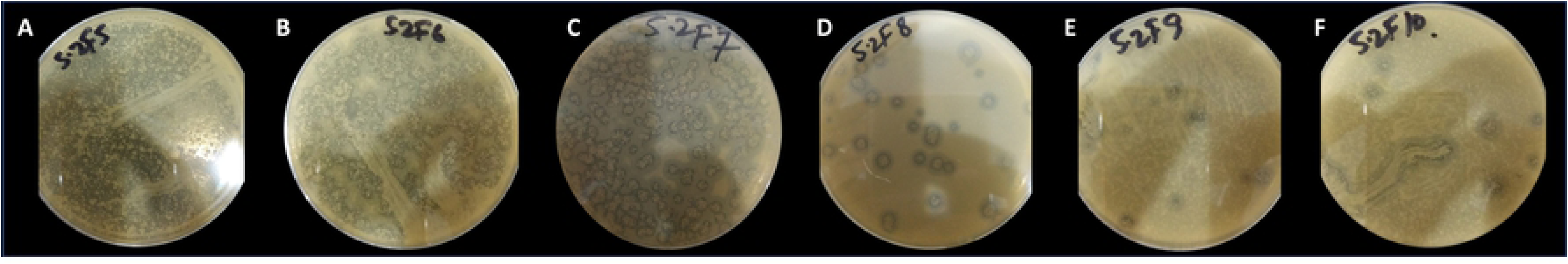
Virus plaques are visible on the lawn of XDR Acinetobacter baumannii strain. Ten plates were prepared but lysis became observable from 10^−5^ dilution and plaques became countable on plate containing 10^−7^ phage dilution. For reference, plate containing 10^−5^ to 10^−10^ dilutions are shown. Note decreasing plaque number from A through F.

### Acinetobacter Clearance Assay

A 0.5 McFarland inoculum of *Acinetobacter baumannii* was prepared in 10 ml tube. Three circles of one-inch diameter each were drawn on a pre-sterilized lab bench surface (Fig 2) and labeled as “phenol”, “phage” and “saline”. Then 2ml of bacterial inoculum was poured in each circle. After 20-25 min, 250 μl of 90% phenol, normal saline, and bacteriophage suspension was added on respective inoculated circles. Phage was added at 10 MOI (multiplicity of infection: ratio between no. of infectious phage particles and no. of host bacterial cells). Same procedure was adopted for two more surfaces; top of the incubator and office table.

After overnight exposure of inoculated circles with phenol (positive control), saline (negative control) and phage, any remaining bacteria were collected with a moistened sterile swab by rolling it over each circle thoroughly. Then cotton part of swab was aseptically cut into 10ml sterile saline and vortexed to collect bacteria in the saline. Subsequently, 1 µl calibrated loop (SPL Life Sciences, Catalog no. 90001) was dipped into the saline containing inoculum from circles and LB agar plates were semi-quantitatively streaked. Colonies from phenol exposed, saline exposed and phage exposed circles were counted to determine the bactericidal effects of phage (Table 1).

**Fig 2:**
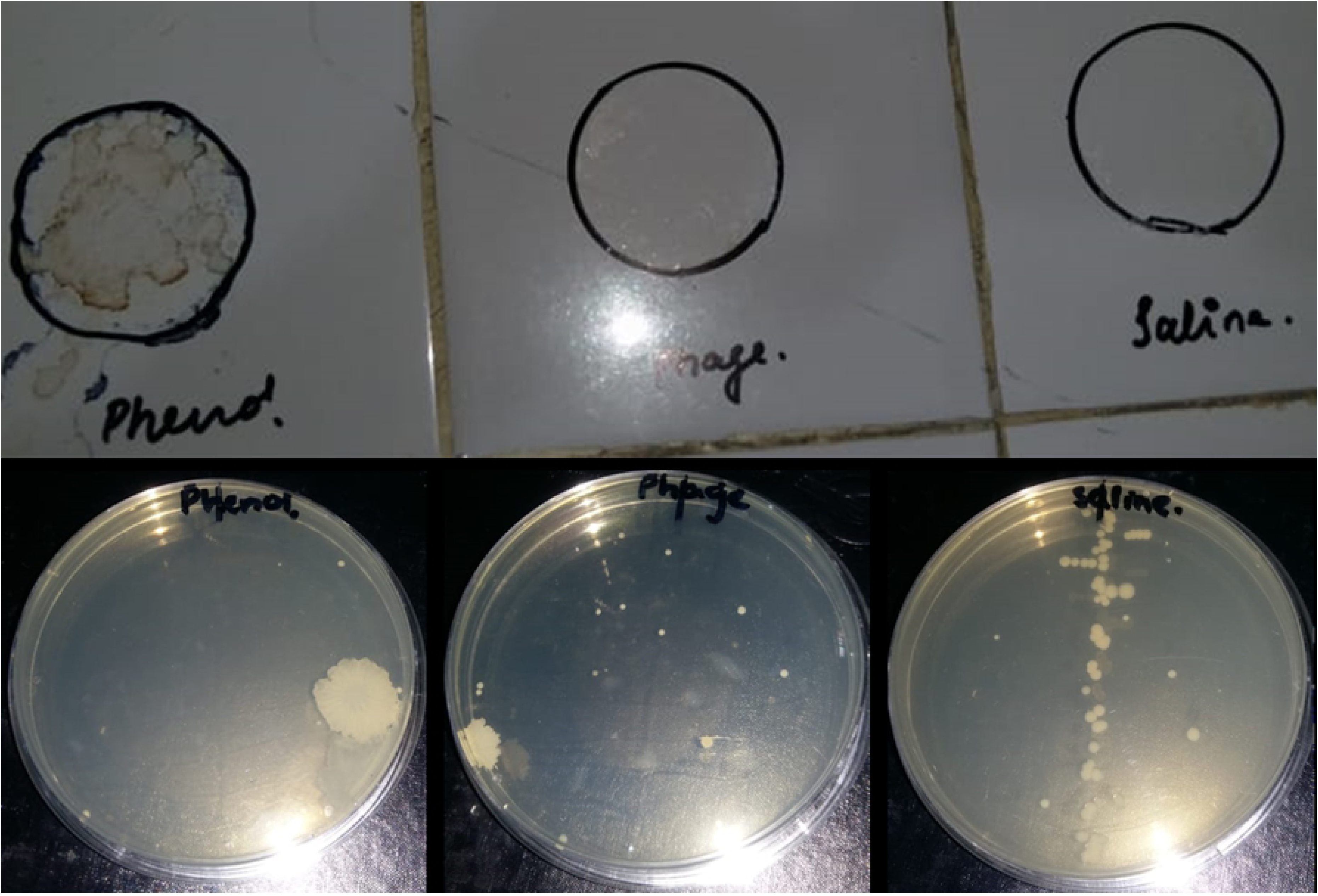
A: Representative illustration of the circled solid surfaces where XDR Acinetobacter baumannii strain was inoculated. Three different surfaces were selected and inoculated as explained in the text and table 1. **B:** Representative plates showing clearance of Acinetobacter from surfaces exposed with either phenol (B1, positive control) or Phage (B2). Plate streaked with saline (B3, negative control) shows much higher growth of bacteria. Note that plates B1 and B2 only have sporadic colonies which do not necessarily represent specific growth originating from the inoculated surfaces.

**Table 1:** Colony forming units (CFU/μL) of Acinetobacter baumannii from phenol, saline and phage exposed surfaces

| Surface | Phenol | Phage | Saline |
| --- | --- | --- | --- |
| Lab Bench | 3 | 15 | 78 |
| Top of Incubator | 5 | 13 | 103 |
| Office Table | 2 | 11 | 91 |
| **Avg±SD** | **3.33±1.5** | **13±2** | **90.67±12.5** |

## Results and Discussion

### Phage Isolation and Titer

We collected some 50 sewage samples from various sites and one sample from Children Hospital Multan yielded bacteriophage. For purification of single phage, a single well isolated plaque was picked from the plate and subjected to enrichment. After every round of enrichment isolated plaque was collected and process was repeated three time in order to get a pure single type of phage (Fig 1). Titer was measured in terms of plaque forming units per milliliter (PFU/ml). There are many ways of measuring virus concentration in a sample. Some of them are based on measuring quantity of physical particles based on viral proteins or antibodies (Such as ELISA and Western blotting). Others measure concentration of virus nucleic acid by qPCR methods. Many particles in a given sample may be incapable of causing productive infection. Most reliable approach, therefore, is measurement of infectious particles because methods based on physical measurements are not always linearly correlated to infectivity [10]. Therefore, PFU/ml gives us infectious titer and hence approximates number of “physically fit” particles. Plates containing 10^−7^ dilution of phage were used to count plaques as it had isolated and hence countable plaques (Fig 1C). All work was done in triplicate and averages were calculated. PFU/ml were extrapolated from the plaque count to be 2×10^12^.

### Use of Phage as Bactericidal Agent

Hospital acquired infections are increasingly becoming a great challenge due to drug resistance and augmented virulence. Several hospital strains hide themselves on solid surfaces such as work benches of hospital staff, side tables of patient beds, and medical instruments such as ventilators [1,4]. Fortunately, more and more health establishments are implementing infection control regimes in their hospitals. Most, however, use phenolics to decontaminate environment due to their low cost, ready availability and great microbicidal potential. But it is not an agent of choice when it comes to its undesired effects. Many studies have shown it to be toxic to animals and humans and especially infants [7]. Its unfriendliness to environment and patients precludes its use in disinfection of medical devices.

In this backdrop, we set out to determine potential of our acinetophage isolate in clearing clinically relevant XDR *Acinetobacter* strain from solid surfaces. For our experiment, we decided to mimic conditions that are common to hospital inhabiting pathogens (Fig 2). We inoculated different solid surfaces with XDR *Acinetobacter baumannii* strain in known quantities and applied our phage on those surfaces to determine its clearing potential. We infected our bacterial inoculums at 10 MOI because calculations based on Poisson distribution model predict that at 10 MOI almost all bacterial cells receive at least one virus particle. Same volumes of 90% phenol and normal saline were used to serve as positive and negative controls respectively. Purpose was to observe effectiveness of phage as compared to phenol which is an established bactericidal.

After overnight exposure to phage, phenol and saline, we collected any remaining bacteria from those surfaces and cultured on LB agar plates in quantitative manner using calibrated 1µl loops. Therefore, colony count represented CFU per microliter after the exposure of our bactericidal agents.

We observed that phenol, quite expectedly, was the most efficient antibacterial as its exposure resulted in least number of CFU from all three surfaces; 3, 5 and 2 from lab bench, incubator top and office table respectively (Table 1). Saline exposed surfaces gave much higher colony count (78, 103 and 91 CFU/µl for three surfaces in above mentioned order). Again, it was an expected observation as saline has no antimicrobial potential. Phage exposed surfaces gave 15, 13 and 11 CFU/µl which, although, was higher number when compared to phenol but is significantly smaller than in case of saline treatment (Table 1).

Colonies appearing from phenol and phage exposed samples were sporadic (Fig 2: B1 and B2). They were not neatly present on the streaked area, rather, they were randomly located on the plates. It can therefore, be concluded that those colonies did not necessarily originate from inoculated surfaces and may have grown from accidental trapping of bacteria from incubator environment. Why, then, we did not see similar number of nonspecific colonies in both phage and phenol exposed inoculums? This apparent anomaly can be explained on the basis of broad-spectrum effect of phenol but not phage. As swabs were used to collect bacteria from inoculated surfaces (as explained in materials and methods), they could have also carried traces of phenol and phages along with bacteria. When these were subsequently cultured on LB plates, phenol and phages (albeit, in small quantities) would have entered in the plates. Any bacteria entering from incubator environment would have been killed by broad-spectrum effect of phenol but not by a narrow-spectrum phage; resulting in more colonies in latter case.

## Conclusion

This experimental assay provides a proof of concept that phages can be used as infection control agents in hospitals. In contrast to phenolics, they have the advantage of being environment friendly, very specific to their bacterial host and completely safe for humans. Their safety is such that they are being investigated systematically for use as human therapeutics worldwide [11,12]. However, their extreme specificity also affords certain limitations. Unlike broad spectrum chemical agents, one type of phage can usually target one bacterial species. Effective broad range phage based bactericidals, therefore, must contain more than one type of phage. Such phage cocktails can be used to kill a spectrum of hospital inhabiting pathogens while being environment, healthcare worker and patient friendly. An effective phage cocktail must contain phages against pathogens implicated in causing hospital acquired infections (e.g. *Acinetobacter baumannii*, *Staphylococcus aureus*, *Pseudomonas aeruginosa*, *Klebsiella pneumoniae* etc.) [5].

